# Hypomethylation in *FASTKD1* detected in the association between *in utero* tobacco exposure and conduct problem in a New Zealand longitudinal study

**DOI:** 10.1101/2021.04.08.438710

**Authors:** Alexandra J. Noble, John F. Pearson, Joseph M. Boden, L. John Horwood, Martin A. Kennedy, Amy J. Osborne

## Abstract

Despite the known adverse effects of *in utero* tobacco exposure on offspring health, maternal tobacco use during pregnancy remains prevalent and is a major driver of health inequalities. One such health inequality is the development of conduct problem (CP) in exposed offspring which may be mediated by methylation changes that persist into adulthood. Here we apply a genome-wide approach to probe the association between maternal tobacco use during pregnancy and CP outcomes in exposed offspring. We examined maternal tobacco use during pregnancy (*in utero* exposure) in the Christchurch Health and Development Study, a longitudinal birth cohort studied for over 40 years. We then evaluated the interaction between methylation effects of *in utero* exposure and CP score. When modelling this interaction between *in utero* exposure and CP score we detected nominal DNA methylation differences, at *FASTKD1* which has roles in early development. Our observations are consistent with DNA methylation mediating the development of CP following *in utero* tobacco exposure. In addition, we detected nominal significance in *FRMDA4* and *MYO1G* between individuals exposed to tobacco *in utero* and those that were unexposed, however these did not reach significance after adjustment for multiple testing. However due to limited power in our analysis, further studies are needed to investigate the interaction between *in utero* tobacco exposure and high CP health outcomes.

## Introduction

The use of tobacco during pregnancy is one of the leading causes of perinatal compromise for developing offspring, and one of the most preventable [1]. For example, low birth weight [2], congenital heart anomalies [3], asthma/respiratory illness [4, 5], and sudden infant death syndrome (SIDS) [6] are all associated with maternal tobacco use during pregnancy, the rate of which remains relatively high in New Zealand (18.4% [7]), despite declining tobacco use rates overall [8].

While immediate perinatal compromise in infants due to maternal smoking is well documented, the long-term effects into later childhood, adolescence and adulthood are not understood. Maternal tobacco use in pregnancy has been associated with later risks of mental health and related adjustment problems in childhood and adolescence [9, 10]. Further, there is also evidence that maternal smoking during pregnancy is associated with increased risks of conduct disorders and antisocial behaviours in offspring [11] [12–14]. This association is not explained by postnatal environment [15]. Further associations have been identified between maternal tobacco use during pregnancy and the increased risk of the development of attention-deficit hyperactivity disorder (ADHD) [16]. Also affected are offspring neurodevelopment and behaviour, suggesting that poor behavioural adjustment (often termed ‘conduct problems’, CP) can be considered a consequence of maternal smoking during pregnancy [11]. While these traits can be linked to other societal risk factors such as low socioeconomic status and early-life adversity [17], their association with maternal tobacco use during pregnancy is intriguing. Tobacco smoking is a strong modifier of DNA methylation [18–20]. We propose that tobacco use during pregnancy could act on CP risk from *in utero* exposure on DNA methylation. Understanding this association is crucial for furthering the paradigm of the developmental origins of human health and disease (DOHaD) [21].

Overall there is a better understanding of the impact of *in utero* tobacco exposure on DNA methylation and only limited preliminary work has been carried out on the interaction between *in utero* tobacco exposure and the onset of CP [22]. Recent research has demonstrated links between prenatal tobacco exposure and specific DNA methylation patterns of newborn offspring [23–26]. Importantly, tobacco-induced DNA methylation changes can persist into adolescence [27] [26, 28].Further, meta-analyses of multiple CpG sites in the gene *GFI1* (Growth Factor Independent one transcriptional repressor) were found to be differentially methylated in adult offspring in response to being exposed to tobacco *in utero* [29]. In contrast, the role of DNA methylation in the association between *in utero* tobacco exposure and CP is less clear. Preliminary work by Sengupta et al. [22] found three loci with modest DNA methylation changes in response to maternal tobacco use during pregnancy and CP phenotypes, but the etiology of this link has not been fully explored. One potential mechanism is that differential DNA methylation induced during the *in utero* period is influencing the development of CP in childhood.

Here we hypothesise that given maternal tobacco use during pregnancy has been previously linked to offspring CP during early childhood and adolescence, and that maternal tobacco use during pregnancy can affect DNA methylation of offspring through to adolescence and adulthood, that DNA methylation is altered at genes that have biological relevance for CP phenotypes, in the whole blood of adults who were exposed to tobacco *in utero*. To test this hypothesis, we used the Illumina EPIC array to interrogate differential methylation in the DNA of participants from the Christchurch Health and Development Study (CHDS) whose mothers consumed tobacco during pregnancy.

## Methods

A cohort of 109 participants from the CHDS (N= 1265 in total) gave blood samples at about age 28 which were used for Illumina EPIC array analysis (Table 1). CHDS participants were chosen based on their *in utero* tobacco exposure status, their adult smoking status, and their CP scores. The *in utero* tobacco exposure was defined as 10+ cigarettes per day throughout pregnancy. A total of 49 individuals fitted this description. Within this group, 23 of the participants were also adult smokers. A total number of 60 individuals were used for the non-exposed control group. Eight of these participants were adult smokers. Furthermore, in both the exposed and non-exposed groups there were individuals who were selected with either a ‘high’ or ‘low’ score on a measure of childhood CP at age 7-9 years. Severity of childhood CP was assessed using an instrument that combined selected items from the Rutter and Conners child behaviour checklists [30–33] as completed by parents and teachers at annual intervals from 7-9 years. Parental and teacher reports were summed and averaged over the three years [34] to derive a robust scale measure of the extent to which the child exhibited conduct disordered/oppositional behaviours (mean (SD)= 50.1 (7.9), range 41-97). For the purposes of this report a ‘high’ score was defined as falling into the top quartile of the score distribution (scores > 53) and a ‘low’ score was defined as < 46.

**Table 1.** Cohort characteristics of the subset of individuals (n = 109) used to assess differential DNA methylation and *in utero* tobacco exposure, and their matched controls, all from the CHDS.

|  | *In utero* tobacco<br>exposed group<br>N= 49 | Non-exposed<br>control group<br>N= 60 |
| --- | --- | --- |
| Sex |  |  |
| Male | 36 | 46 |
| Female | 13 | 14 |
| Paternal<br>socioeconomic status |  |  |
| 1 | 3 | 13 |
| 2 | 21 | 30 |
| 3 | 25 | 17 |
| Adult tobacco smoking<br>status |  |  |
| Never smoker | 26 | 52 |
| Regular<br>smoker | 23 | 8 |
| Conduct problem<br>score (CP) |  |  |
| Low CP (< 46) | 26 | 41 |
| High CP (> 53) | 23 | 19 |

All samples taken were whole blood and collected at approximately 28 years old, and DNA extractions were conducted using the Kingfisher Flex System (Thermo Scientific, Waltham, MA USA), as per the published protocols. DNA was quantified via NanoDrop™ (Thermo Scientific, Waltham, MA USA) and standardised to 100ng/μl. Equimolar amounts were shipped to the Australian Genomics Research Facility (AGRF, Melbourne, VIC, Australia) for processing via the Infinium® Methylation EPIC BeadChip (Illumina, San Diego, CA USA). The arrays were carried out in groups over three batches. One in 2016, one in 2017 and another in 2020. Analysis was carried out in R statistical software (Version 3.5.2). Quality control checks were performed on the raw data, firstly sex chromosomes and a total of 90 failed probes (detection P value of < 0.01 in at least 50% of samples) were excluded from the analysis. CpG sites known to be problematic due to adjacent single nucleotide variants or which did not map to a location in the genome were also excluded [35]. This left a total of 699,916 CpG sites for further analysis. Pre-processing was performed using Noob [36]. Normalisation was then visually inspected for performance using beta density distribution plots and Multi-dimensional scaling of the 5,000 most variable CpG sites. Hierarchical regression was used to investigate the best linear model to be fitted to the methylated/unmethylated or M ratios. The two following models were picked as the most robust equations to assess for differential DNA methylation for our data set.

### *In utero* exposed only (model 1)

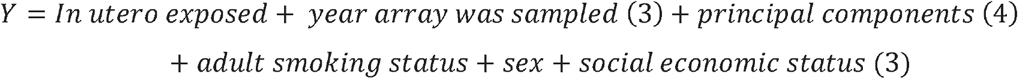

### *In utero* exposed with the interaction of CP (model 2)

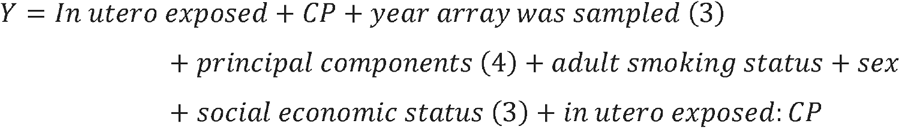

Analysis corrected for the following variables: i) year array was sampled (3 levels), and; ii) population stratification (four principal components from 5000 most variable SNPs [37]), adult tobacco status (bivariate), sex (bivariate), socioeconomic status (three levels). Q-Q plots of the residuals were also used to generate lambda values to assess for over-inflation. Linear regression models were used to generate the top tables of differentially methylated CpG sites and were corrected for multiple testing using the Benjamini-Hochberg method (BH). Differentially methylated CpG sites that were intergenic were matched to the nearest neighbouring genes in Hg19 using the R package GenomicRanges [38]. The package ggplot (Version 3.3.2) was used to construct all dotplot graphs [39], with the log transformed normalised methylated and unmethylated values plotted.

## Results

### Data analysis

Both the *in utero* exposed only (model 1), and the *in utero* exposed with the interaction of CP (model 2) were assessed for genomic inflation via Q-Q plots (Supplementary Figure 1A and B). No significant indication of inflation was observed for either of the two models: *in utero* exposed only with λ = 0.97 (Supplementary Figure 1A) and *in utero* exposed with the interaction of CP with λ = 1.03 (Supplementary Figure 1B).

### Methylation differences from *in utero* exposed only individuals compared to non-exposed controls (model 1)

Firstly, differences in DNA methylation in response to *in utero* exposure were assessed, while adjusting for year array was sampled, four principal components, adult tobacco smoking status, sex, and socioeconomic status. Results of this analysis identified nominally significant differential DNA methylation between individuals exposed to tobacco *in utero* compared to the non-exposed control group. A total of 653 CpG sites had a P value less than 0.001, of these 222 were hypomethylated and the rest (431) were hypermethylated.

Top tables of the most significant CpG sites were then constructed; the top 30 CpG sites are displayed in Table 2. Several nominally significant CpG sites were observed, although none were significant after adjustment for multiple testing. The top five differentially methylated sites resided in four genes: two sites in *FRMD4A* (5.1% and 4.6% differentially methylated), and one each in *MYOG1* (7.4% differential methylation), *WIPF3* (3.1% differential methylation) and *RTN1* (3.1% differential methylation). At all five of these CpG sites, differences decreased, indicating hypomethylation in response to *in utero* tobacco exposure. This same trend is observed in the other remaining top CpG sites, as 28/30 were hypomethylated but not significant in the exposed group. This is further illustrated by scatter plots of the top four most significant CpG sites in Figure 1.

**Table 2.** Top differentially methylated CpG sites to *in utero* tobacco exposure in offspring. Beta values with P values, nominal and adjusted by the Benjamini and Hochberg method. Locations are relative to hg19 with gene names for overlapping genes or nearest 5□.

| Illumina ID | Gene | Chromosome | Non exposed | Exposed | Difference | P value | Adjusted P value |
| --- | --- | --- | --- | --- | --- | --- | --- |
| cg25464840 | *FRMD4A* | 10 | 0.708 | 0.755 | 0.047 | 6.14E-07 | 0.247 |
| cg15507334 | *FRMD4A* | 10 | 0.598 | 0.649 | 0.051 | 7.06E-07 | 0.247 |
| cg04180046 | *MYO1G* | 7 | 0.513 | 0.587 | 0.074 | 1.45E-06 | 0.337 |
| cg20745684 | *WIPF3* | 7 | 0.254 | 0.285 | 0.031 | 2.04E-06 | 0.356 |
| cg01604380 | *RTN1* | 14 | 0.218 | 0.250 | 0.032 | 2.67E-06 | 0.374 |
| cg22931725 | *FMN2* | 1 | 0.051 | 0.059 | 0.008 | 3.75E-06 | 0.391 |
| cg06284231 | *CLEC14A* | 14 | 0.161 | 0.183 | 0.023 | 4.07E-06 | 0.391 |
| cg02721176 | *C10orf96* | 10 | 0.300 | 0.363 | 0.064 | 4.95E-06 | 0.391 |
| cg02841155 | *SLC27A6* | 5 | 0.257 | 0.304 | 0.047 | 5.37E-06 | 0.391 |
| cg27339941 |  | 9 | 0.562 | 0.600 | 0.038 | 5.59E-06 | 0.391 |
| cg01026094 |  | 20 | 0.284 | 0.322 | 0.038 | 6.36E-06 | 0.405 |
| cg24601030 | *DIXDC1* | 11 | 0.756 | 0.797 | 0.041 | 8.04E-06 | 0.469 |
| cg15433297 | *PKHD1L1* | 8 | 0.322 | 0.346 | 0.024 | 8.95E-06 | 0.482 |
| cg05798339 |  | 3 | 0.344 | 0.359 | 0.015 | 1.13E-05 | 0.566 |
| cg21771773 |  | 5 | 0.164 | 0.201 | 0.037 | 1.21E-05 | 0.566 |
| cg15766464 |  | 5 | 0.477 | 0.505 | 0.029 | 1.37E-05 | 0.597 |
| cg20595752 | *TEAD3* | 6 | 0.524 | 0.559 | 0.036 | 1.48E-05 | 0.610 |
| cg06926300 |  | 10 | 0.159 | 0.168 | 0.008 | 1.64E-05 | 0.638 |
| cg11813497 | *FRMD4A* | 10 | 0.767 | 0.817 | 0.050 | 1.87E-05 | 0.666 |
| cg24149528 |  | 15 | 0.349 | 0.385 | 0.036 | 1.90E-05 | 0.666 |
| cg10068883 | *FAM166B* | 9 | 0.595 | 0.637 | 0.043 | 2.10E-05 | 0.693 |
| cg11157260 |  | 4 | 0.738 | 0.767 | 0.029 | 2.31E-05 | 0.693 |
| cg13206397 |  | 16 | 0.891 | 0.881 | -0.010 | 2.37E-05 | 0.693 |
| cg16081285 | *CSRNP3* | 2 | 0.139 | 0.159 | 0.020 | 2.37E-05 | 0.693 |
| cg19453818 | *CAV2* | 7 | 0.408 | 0.438 | 0.030 | 2.70E-05 | 0.726 |
| cg15688339 |  | 5 | 0.384 | 0.446 | 0.062 | 2.83E-05 | 0.726 |
| cg03770646 |  | 19 | 0.640 | 0.677 | 0.038 | 3.07E-05 | 0.726 |
| cg06372850 |  | 2 | 0.222 | 0.265 | 0.043 | 3.07E-05 | 0.726 |
| cg14644418 | *FOXI2* | 10 | 0.824 | 0.852 | 0.028 | 3.23E-05 | 0.726 |
| cg25072920 | *TCN1* | 11 | 0.944 | 0.940 | -0.004 | 3.39E-05 | 0.726 |

**Figure 1.**
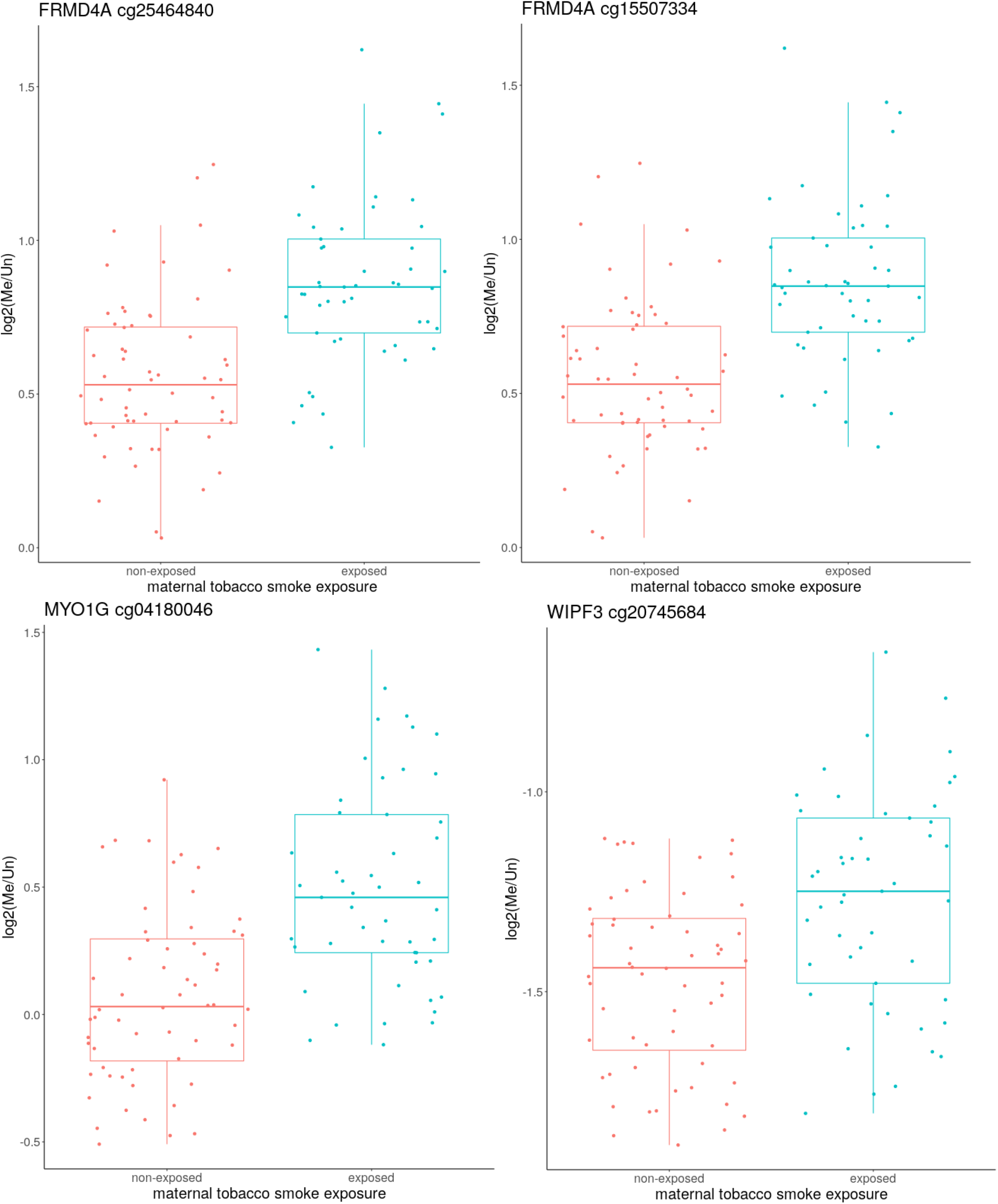
The top four CpG sites differentially methylated due to *in utero* maternal tobacco exposure, these sites resided in genes *FRMD4A, MYO1G* and *WIPF3*. The log transformed normalised beta values of the non-exposed individuals are seen on the left, with the exposed *in utero* to tobacco individuals on the right.

### Methylation differences between *in utero* exposed with the interaction of conduct problems (model 2)

To investigate genome-wide differential methylation specific to the interaction between *in utero* tobacco exposure and CP score, we determined differential methylation between i) individuals with high CP vs. low CP scores who were exposed to tobacco *in utero*, and; ii) individuals with high or low CP scores who were not exposed to tobacco *in utero)*. Dividing the data in this way allowed us to specifically ask whether exposure to tobacco *in utero* may be associated with a high CP score as a result of an alteration to DNA methylation.

Top tables of the most significantly differentially methylated CpG sites were constructed (model 2), with the top 30 CpG sites displayed in Table 3. No CpG sites were found to be significant after adjustment for multiple testing. Within the top 30 CpG sites, an even split was seen between hypo and hypermethylation. The magnitude of the differential methylation in the *in utero* exposed low vs. high CP, and the non-exposed low vs. high CP under the interaction model are all less than 1.2%. The top four most differentially methylated sites under this interaction are plotted in Figure 2 (CpG site cg12163448 in *FASTKD1*, cg01210554 with no known associated gene, cg13339919 in *SLC10A7* and cg21516989 in *LPIN1* on individual plots). Scatter plots, overlaid with a box, are fitted to the four subcategories for each CpG site: i) non-exposed low CP; ii) non-exposed high CP; iii) *in utero* exposed low CP, and; iv) *in utero* exposed high CP. Each of the four subcategories here has a range of methylation values for the individuals within the group.

**Table 3.** Top differentially methylated CpG sites between *in utero* maternal tobacco exposure and the interaction with CP. Beta values with P values, nominal and adjusted by the Benjamini and Hochberg method. Locations relative to hg19 with gene names for overlapping genes or nearest 5□ gene.

| Illumina ID | Gene | Chromosome | *In utero*<br>maternal<br>tobacco<br>exposed<br>low CP | *In utero*<br>maternal<br>tobacco<br>exposed<br>high CP | Difference | P value | Adjusted P<br>value |
| --- | --- | --- | --- | --- | --- | --- | --- |
| cg12163448 | *FASTKD1* | 2 | 0.142 | 0.155 | 0.013 | 1.01E-06 | 0.459 |
| cg01210554 |  | 7 | 0.060 | 0.061 | 0.001 | 1.31E-06 | 0.459 |
| cg13339919 | *SLC10A7* | 4 | 0.9457 | 0.9456 | -0.0001 | 2.08E-06 | 0.486 |
| cg21516989 | *LPIN1* | 2 | 0.946 | 0.947 | 0.001 | 6.89E-06 | 0.962 |
| cg17343033 |  | 5 | 0.881 | 0.882 | 0.001 | 8.08E-06 | 0.962 |
| cg01394525 | *LAMC3* | 9 | 0.910 | 0.915 | 0.005 | 8.82E-06 | 0.962 |
| cg02809796 | *PCGF3* | 4 | 0.9185 | 0.9186 | 0.0001 | 9.62E-06 | 0.962 |
| cg12361925 | *FAM134A* | 2 | 0.043 | 0.042 | -0.001 | 1.37E-05 | 0.984 |
| cg23048036 | *CEP97* | 3 | 0.808 | 0.796 | -0.012 | 1.71E-05 | 0.984 |
| cg05750962 |  | 18 | 0.082 | 0.083 | 0.001 | 2.80E-05 | 0.984 |
| cg24835473 | *CHCHD6* | 3 | 0.931 | 0.929 | -0.002 | 3.33E-05 | 0.984 |
| cg23399286 | *OC90* | 8 | 0.8087 | 0.8088 | 0.0001 | 3.81E-05 | 0.984 |
| cg16546489 | *HSD17B1* | 17 | 0.918 | 0.915 | -0.003 | 3.84E-05 | 0.984 |
| cg09692779 |  | 15 | 0.923 | 0.921 | -0.003 | 3.88E-05 | 0.984 |
| cg14496346 |  | 9 | 0.095 | 0.094 | -0.001 | 4.02E-05 | 0.984 |
| cg14420953 |  | 6 | 0.938 | 0.935 | -0.003 | 4.10E-05 | 0.984 |
| cg09125477 | *C16orf91* | 16 | 0.102 | 0.111 | 0.009 | 4.11E-05 | 0.984 |
| cg11095011 |  | 6 | 0.936 | 0.935 | -0.001 | 4.27E-05 | 0.984 |
| cg24753210 | *PPP1R14B* | 11 | 0.068 | 0.066 | -0.001 | 4.83E-05 | 0.984 |
| cg25849390 | *CCT6A* | 7 | 0.884 | 0.880 | -0.004 | 5.21E-05 | 0.984 |
| cg13787134 | *PHF2* | 9 | 0.9391 | 0.9392 | 0.0001 | 5.46E-05 | 0.984 |
| cg05858227 |  | 13 | 0.947 | 0.945 | -0.002 | 5.58E-05 | 0.984 |
| cg18168310 |  | 11 | 0.672 | 0.686 | 0.014 | 5.69E-05 | 0.984 |
| cg06923651 | *SPINK4* | 9 | 0.437 | 0.438 | 0.001 | 6.06E-05 | 0.984 |
| cg04807567 | *MYO3B* | 2 | 0.850 | 0.848 | -0.002 | 6.25E-05 | 0.984 |
| cg03243165 |  | 2 | 0.9393 | 0.9398 | 0.0005 | 6.45E-05 | 0.984 |
| cg03492068 |  | 17 | 0.9100 | 0.9101 | 0.0001 | 7.06E-05 | 0.984 |
| cg10671167 | *CDK13* | 7 | 0.083 | 0.084 | 0.001 | 7.47E-05 | 0.984 |
| cg14778074 | *GABRA2* | 4 | 0.110 | 0.108 | -0.002 | 7.60E-05 | 0.984 |
| cg09482386 | *CYB561* | 17 | 0.941 | 0.940 | -0.002 | 8.47E-05 | 0.984 |

**Figure 2.**
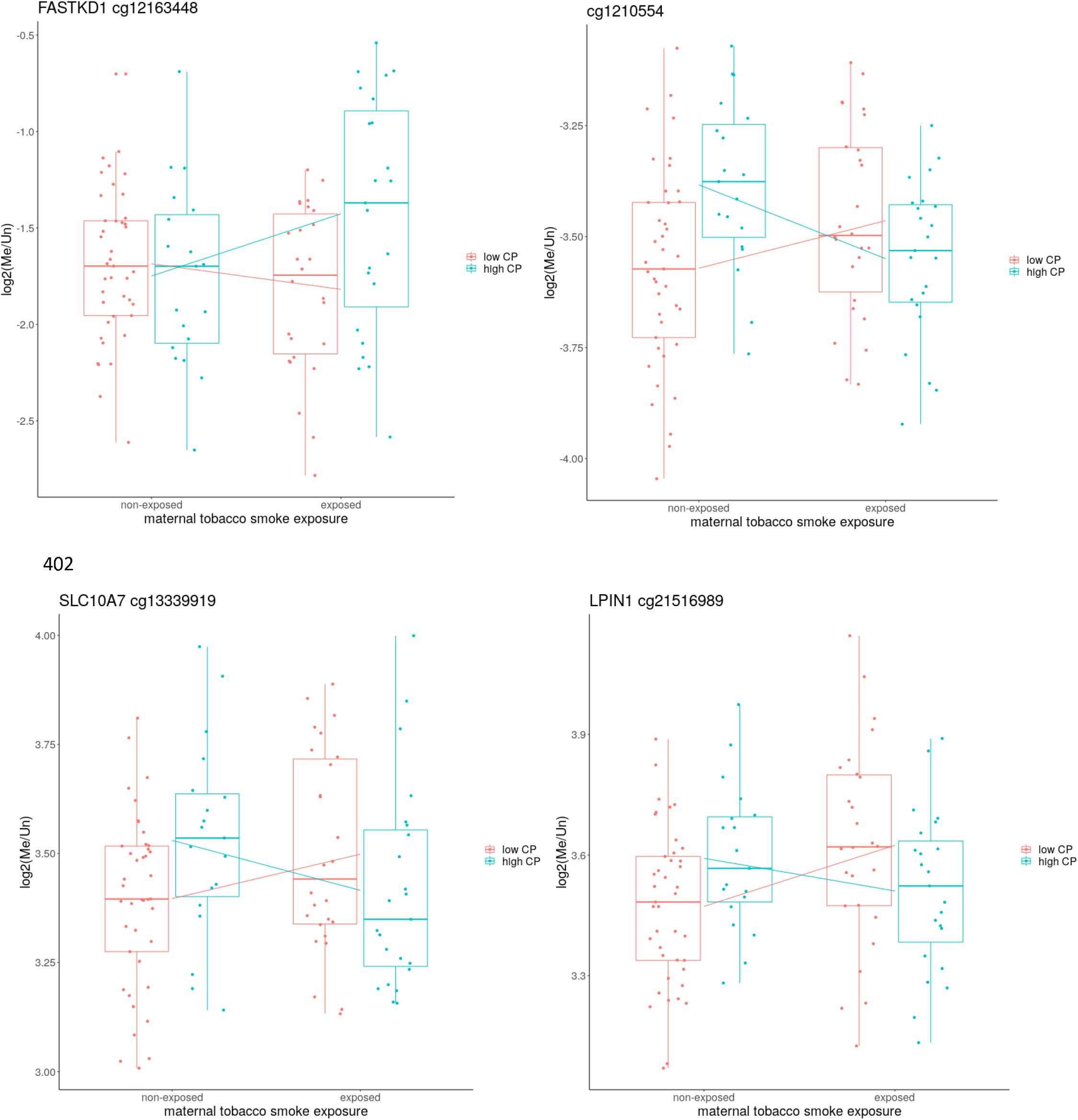
The top four most significantly differentially methylated (nominal P < 0.01) CpG sites when *in utero* tobacco exposure was assessed with the interaction CP. Non-exposed individuals are plotted on the left of each plot, colour coded for either low CP (salmon) or high CP (cyan), with exposed individuals on the right. Lines from the non-exposed group to the exposed group represent the difference in average methylation between non-exposed and exposed with (salmon) and without (cyan) CP.

## Discussion

The effect of *in utero* tobacco exposure has been widely found to induce differential DNA methylation marks in the genomes of exposed offspring, however much of what we know comes from studies involving infants and young children. Here, we investigated the effect of *in utero* tobacco exposure on DNA methylation in the blood of adult New Zealanders who were exposed to tobacco during development and explored the association between exposure during development with a high conduct problem score in childhood and adolescence.

Firstly, we assessed the effect of exposure to tobacco *in utero* on offspring DNA methylation, compared to non-exposed controls, without the interaction of CP score. No CpG sites reached an adjusted P value of significance (Table 2). However, the top nominally significant CpG sites were found in the genes *FRMD4A, MYO1G, WIPF3*, and *RTN1*. CpG sites in genes *MYO1G* and *FRMD4A* have been previously found to be differentially methylated in response to maternal smoking during pregnancy [40–42]. However, the CpG sites cg20745684 and cg01604380 within the genes *WIPF3* and *RTN1* respectively have not been implicated in any previous study.

Myosin 1G (*MYO1G*) and FERM Domain Containing 4A (*FRMD4A*) are more established biomarkers for *in utero* tobacco exposure [40–42]. However, these prior results come from the analysis of DNA methylation during childhood. Here, we show that methylation differences at these well-established biomarkers, in response to *in utero* tobacco exposure, are detectable in exposed individuals through to adulthood (age ~28 years), therefore our results expand on this current knowledge by demonstrating a potential for long-term stability of developmentally-induced methylation. We suggest that *MYO1G* and *FRMD4A* are specifically differentially methylated in response to maternal tobacco use during pregnancy, and that these methylation changes induced *in utero* appear to be stable into adulthood. Knowledge of the longevity of potentially developmentally-induced methylation marks may be important for understanding later-life disease risk, particularly given the association between *MYO1G* and immunity [43] and *FRMD4A* [44] with Alzheimer’s disease.

Secondly, we assessed the interaction between *in utero* tobacco exposure and CP in exposed offspring on DNA methylation. Thus far, limited evidence has provided a molecular mechanism of the known association between maternal tobacco use during pregnancy and the development of CP in exposed offspring. While no CpG sites displayed genome-wide significance under this interaction model, nominal significance was observed. Beta differences within the top 30 most nominally (P< 0.01) significant CpG sites were small, with the largest being 1.2%. The most differentially methylated CpG sites resided in the following genes: *FASTKD1, SLC10A7* and *LP1N1*.

Methylation at the CpG site in FAST Kinase domain 1 (*FASTKD1*) was elevated in those with high CP scores only in the exposed group (Figure 2). No methylation differences were found in those not exposed to maternal tobacco. Thus, this site in FASTKD1 is a possible mediator of the known effect of maternal tobacco on CP. *FASTKD1* plays a role in the regulation of mitochondrial RNA [45, 46]. Single nucleotide variants within this gene have been associated with glaucoma [47], furthermore, this is evidence to suggest that children with vision impairment are more likely to be diagnosed with disorders such as ADHD than those that do not [48].

Solute Carrier Family 10 Member 7 (*SLC10A7)*, Lipin 1 (*LPIN1*) and Laminin gamma 3 (*LAMC3*) also contained top differentially methylated CpG sites. However, the interaction here is different to what was observed in *FASTKD1*. Differences in methylation are seen in the unexposed CP group. Thus, we hypothesise that these sites may be driven more so by unexposed high CP when compared to the exposed groups rather than the interaction of only exposed high CP. Furthermore, these genes do have functional relevance to the high CP phenotype. Single nucleotide variances in *LIPIN1* have been associated with neurofibrillary tangles which is a pathological protein aggregate found in Alzheimer’s disease [49]. *LAMC3* has diverse roles in cell migration, apoptosis and adhesion, and variants within this gene have been found to contribute to cortical malformations [50–52]. It has been reported that individuals with these variants also have specific behavioural outcomes, e.g., impairments in endogenous attentional processes [51], which collectively suggests that these data may have biological relevance to the CP phenotype.

Therefore, while we detect biologically relevant nominally significant interactions between DNA methylation at individual CpG sites and high CP score in individuals exposed to tobacco *in utero*, further work is required to better characterise the precise mechanism of CP development. CP is a highly complex phenomenon that encompasses a range of phenotypes [53], which will have a range of aetiologies. Hence the mechanism of CP development for non-exposed individuals with high CP is likely to be different to the individuals who were exposed to maternal tobacco smoke with the same phenotype. Our data identify potential epigenetic changes that could mediate the association between maternal smoking and conduct problems which persist into adulthood. These candidates require confirmation in larger independent studies.

## Conclusion

Our findings here suggest that DNA methylation could be involved in the link between *in utero* tobacco exposure and conduct problems. While we were unable to identify genome wide differences in our cohort (N= 109), our results indicate that it would be beneficial to explore our nominal associations in a larger cohort. Further findings may help to elucidate the detrimental effect that *in utero* tobacco exposure has on the genome of exposed individuals, and suggest that disease associated DNA methylation which occurs early in the life course persists into adulthood.

## Ethics approval and consent to participate

All aspects of the study were approved by the Southern Health and Disability Ethics Committee, under application number CTB/04/11/234/AM10 “Collection of DNA in the Christchurch Health and Development Study”.

## Funding

Seed grant - College of Science New Ideas Seed Grant to AJO.

CHDS was funded by the Health Research Council of New Zealand (Programme Grant 16/600)

CMRF provided the funding to write up the findings of this study.

---

**Supplementary Figure 1.**
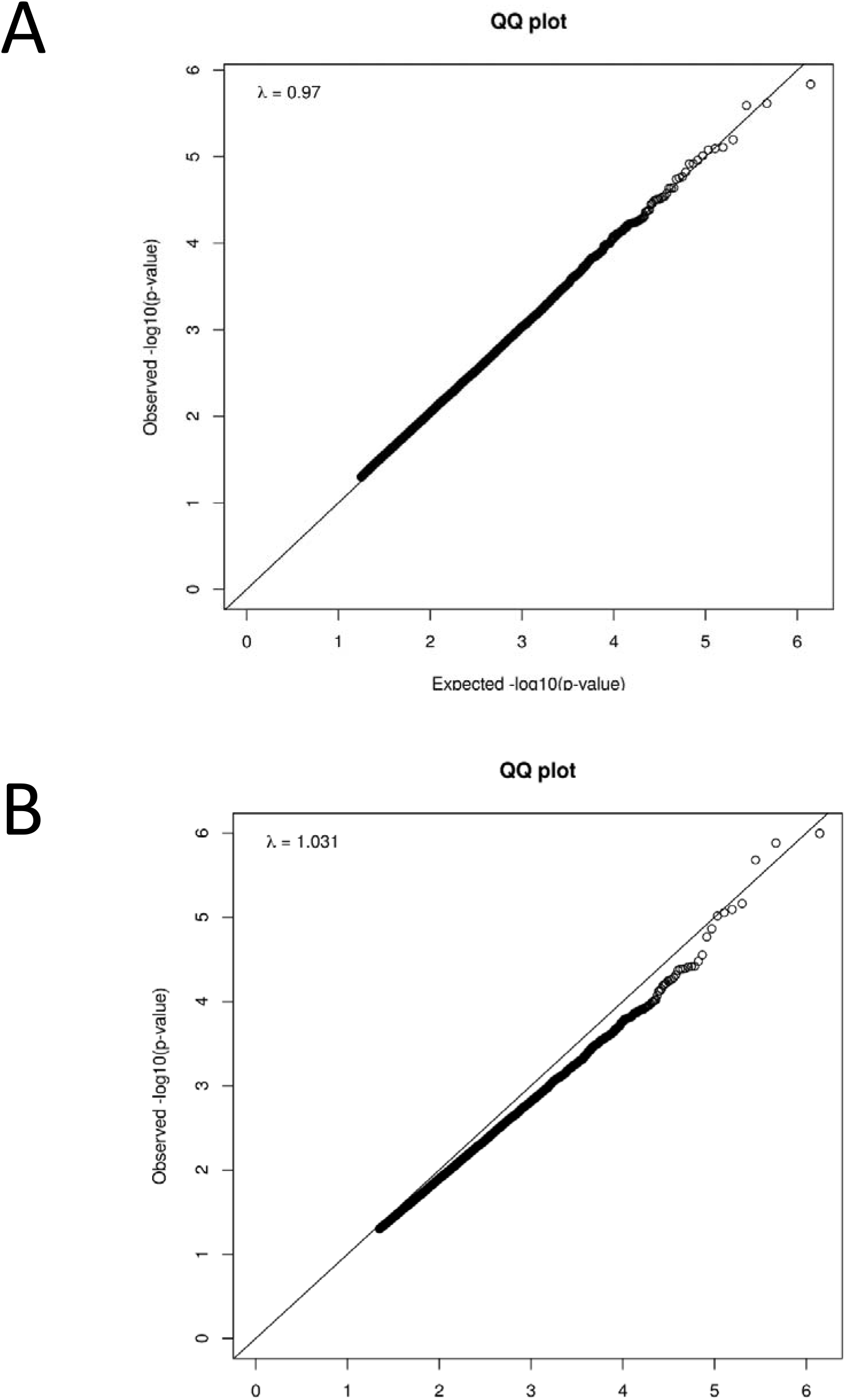

## Notes

### Competing Interest Statement

The authors have declared no competing interest.

